# Learning fly pose from vision-language annotations

**DOI:** 10.64898/2026.09.30.755712

**Authors:** Joshua W. Shaevitz, Matthew G. Sierra, Sarah Kuo Kim, Scott D. Pletcher

## Abstract

Animal pose estimation typically requires manual annotation of body parts to train neural networks for each new recording system or environmental condition. Here we use a general-purpose vision-language model (VLM) to generate annotations in multi-animal recordings of fruit flies (*Drosophila melanogaster*). Initial annotations follow written anatomical descriptions and are reviewed by an independent VLM session using neighboring video frames. On 120 held-out fly images, AI annotations had a median distance of 2.28 pixels from human labels, with 84.0% agreement within eight pixels, averaged across landmark types. The comparison included 10 landmarks, with head and eyes analyzed separately because of systematic differences in placement. Review improved agreement primarily at the feet, with the largest remaining discrepancies at positions with limited visual evidence. Visual inspection revealed cases in which the AI annotations were more anatomically accurate than those of an experienced human annotator, particularly at the feet. A convolutional network trained using SLEAP on 300 AI-annotated fly images showed similar agreement with human labels and closely reproduced the AI annotations on held-out images, with a median distance of 1.56 pixels. We also applied the procedure to recordings from another laboratory, using new AI annotations to train a separate pose network. Although VLM annotation required minutes per fly, the trained networks enabled pose estimation throughout more than 700,000 frames. These results demonstrate that VLM annotations can support fast animal pose estimation without manually labeled training images.

## 1 Introduction

Measurements of coordinated body-part motion connect animal behavior to its neural and biomechanical control [1–3]. Convolutional neural network (CNN)-based tools such as DeepLabCut, DeepPoseKit, LEAP and SLEAP estimate these motions by learning anatomical landmark positions from human-annotated images [4–7]. Their ability to learn from small training sets has made pose estimation applicable to many experimental systems, although each new combination of animal, camera and environment generally requires additional labeling. For example, in the original LEAP study, manually annotating 100 single-fly images with a 32-point skeleton required several hours [5]. Most scientific applications require substantially larger labeled datasets, and multiple animals and occluded appendages further complicate the assignment of anatomical identities and positions.

Several sources of information can reduce the burden of annotation. Semi-supervised methods combine labeled examples with spatial and temporal structure in unlabeled video [8–10], and pretrained animal-pose models can transfer information across datasets [11]. Other methods discover keypoints through reconstruction or prediction [12–15] or introduce anatomical structure through pose priors [16]. Synthetic images can provide known landmark positions from geometric animal models, including models of fruit flies [17–19]. These approaches differ in how they specify the anatomical coordinates to be learned.

Vision-language models (VLMs) offer a means of specifying those coordinates through written descriptions of anatomy. Language-guided pose models associate named structures with image locations [20–22], and general-purpose VLMs can supply annotations for a separate prediction model [23]. The use of a large model to supervise a smaller network is related to model distillation, in which a smaller student model learns to reproduce the outputs of a larger teacher [24]. VLM-derived semantic supervision has previously been incorporated into human-pose training [25], and BehaviorVLM has demonstrated direct anatomical annotation using fluorescent landmarks, human-labeled seed examples, temporal information and multiple camera views [26].

Annotating pose in single-camera recordings requires identifying each appendage and locating its landmarks even when structures overlap or disappear from view. This is particularly challenging in fruit flies, whose six articulated legs frequently cross or pass beneath the body and wings. Movement across adjacent frames can help resolve these ambiguities by revealing which leg a visible segment belongs to and constraining the position of an obscured endpoint in the frame being labeled. A reasoning model can combine this temporal context with anatomical descriptions to assign landmark identities and coordinates in the target frame.

Here we show that visual reasoning can supply the anatomical annotations needed to train fly-pose networks without manually labeled training images. We combine VLM annotation and temporal review with fast convolutional pose estimation and apply the procedure independently to two datasets acquired with different imaging setups. Comparisons with human annotations test both the VLM labels and the resulting network predictions, and a separate experiment in another laboratory demonstrates the procedure under different recording conditions. Together, these experiments establish a route from written descriptions of anatomy to automated pose measurement. The accompanying resources are available through SIESTA (Self-supervised Inference Engine for Spatial Tracking of Animals).

## 2 Results

### 2.1 Learning pose from visual reasoning

We recorded 16 fruit flies (Oregon-R *Drosophila melanogaster*) in four separate arenas, with four flies per arena and all four arenas imaged simultaneously in a single field of view (Fig. 1A). We acquired two recordings of approximately one hour each, imaging the flies from below at 1536 × 1536 pixels and 100 frames per second. Human annotations were performed on 7,083 fly instances in 447 frames. We reserved these frames for comparing VLM annotations and CNN predictions with the human labels. To construct the CNN training set, we used the VLM to annotate temporally separated images from the same recordings (Methods).

**Figure 1:**
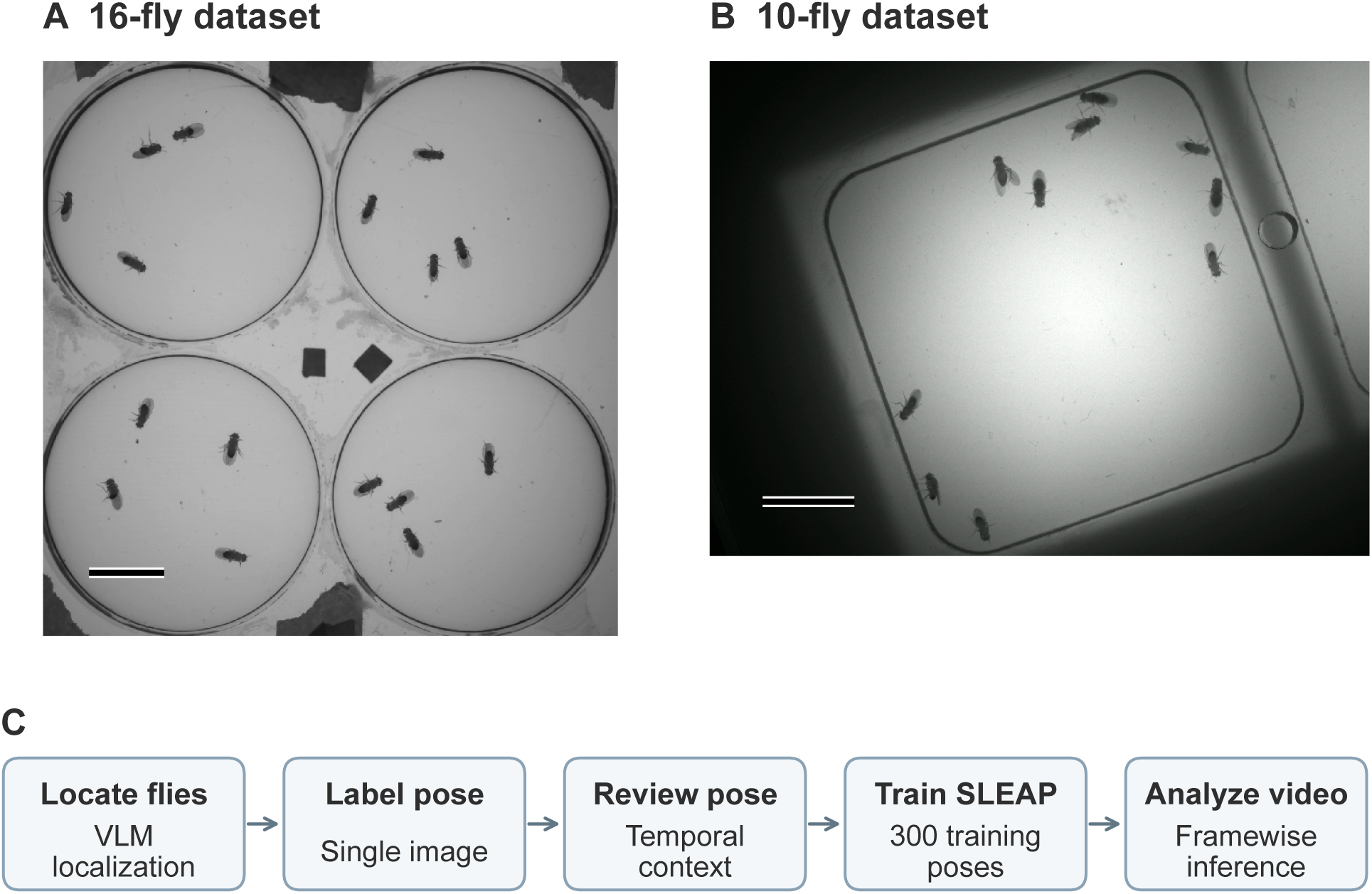
Recording systems and VLM-supervised pose estimation. (A) Single movie frame showing four separate arenas, each containing four flies, imaged simultaneously. (B) A separate experiment with ten flies sharing a single larger arena, recorded with a different imaging setup in another laboratory. Full-frame dimensions are 1536 × 1536 and 1440 × 1080 pixels, respectively; scale bars, 200 pixels. (C) A VLM localizes flies and proposes 13 landmarks for each sampled crop. An independent VLM review inspects the pose and selected temporally adjacent video frames. AI-reviewed poses train the body-orientation and final pose networks, while separate VLM localization annotations train the detector.

The procedure began by locating flies in the full video frames (Fig. 1C). A VLM supplied localization annotations for 100 frames, of which 80 were used to train a detector and 20 for validation. We applied this detector to a pool of 500 frames to generate crops centered on individual flies. A VLM screened the crops for viewpoint, overlap, completeness and broad leg posture before we selected 300 training instances from 150 source frames and 60 validation instances from 60 other frames for pose annotation.

Pose annotation proceeded in two passes. In the first pass, GPT-6 Astra inspected each crop and an enlarged view of the same image, using written anatomical descriptions to identify the head, thorax and abdomen, two eye positions, two wing tips and six terminal foot positions. It returned coordinates for all 13 landmarks, together with descriptions of the anatomical connections along each leg. Each position was classified as directly observed, inferred from partial visual evidence, or weakly constrained by the available visual evidence.

In the second pass, an independent VLM session inspected the target image, the proposed pose and its supporting descriptions. The reviewer first identified uncertainties about landmark position or anatomical identity, then requested temporally adjacent frames to address those uncertainties. It was allowed to inspect up to three batches of eight images within ±0.5 s of the target, choosing subsequent views based on what earlier views revealed. The reviewer used changes in articulation and visibility to reassess landmark positions in the target frame, revising coordinates when warranted and retaining uncertainty where the evidence remained inconclusive.

The AI-reviewed annotations then supervised the body-orientation and final pose networks. We first trained a network on the body and eye landmarks to estimate the head–thorax axis, which we used to rotate crops and landmark coordinates into a common orientation. The aligned 13-point poses supervised a SLEAP U-Net trained from random initialization [7, 27]. Training comprised 37,500 minibatch updates, with checkpoint selection based on the 60 VLM-labeled validation instances. For subsequent video analysis, the detector located flies, the body-orientation network aligned their crops, and the pose network estimated the 13 landmark positions from each image. We then transformed these predictions back into the original video coordinates.

### 2.2 VLM annotations compared with human annotations

To evaluate the VLM annotations, we obtained initial and AI-reviewed poses for a separate set of 120 test flies, with 60 from each recording. These instances were selected from the fixed detector output, and all 120 could be matched to human-annotated animals. Head and eye labels differed systematically in their placement between the human and VLM annotations, as discussed below. We first report the main agreement measurements for the remaining 10 landmarks: thorax, abdominal tip, two wing tips and six feet, comprising 1,199 eligible positions. A complete 13-landmark analysis is presented in Appendix B.

The VLM annotations generally agreed closely with the human labels, with a median distance of 2.28 pixels (Fig. 2A). Averaging across landmark types, 84.0% of positions were within eight pixels of the human reference (Table 1). We find a substantial difference between body parts. Agreement at the same threshold exceeded 98% for the thorax, abdominal tip and wings, but averaged 74.1% for the feet which are much smaller and harder to locate for both AI and human (Fig. 2B). Many foot annotations were as close to the human labels as those of other structures, with a median distance of 2.13 pixels. Their distance distribution, however, had a much broader tail. The 90th percentile reached 21.22 pixels for feet but remained below five pixels for body and wing landmarks.

**Figure 2:**
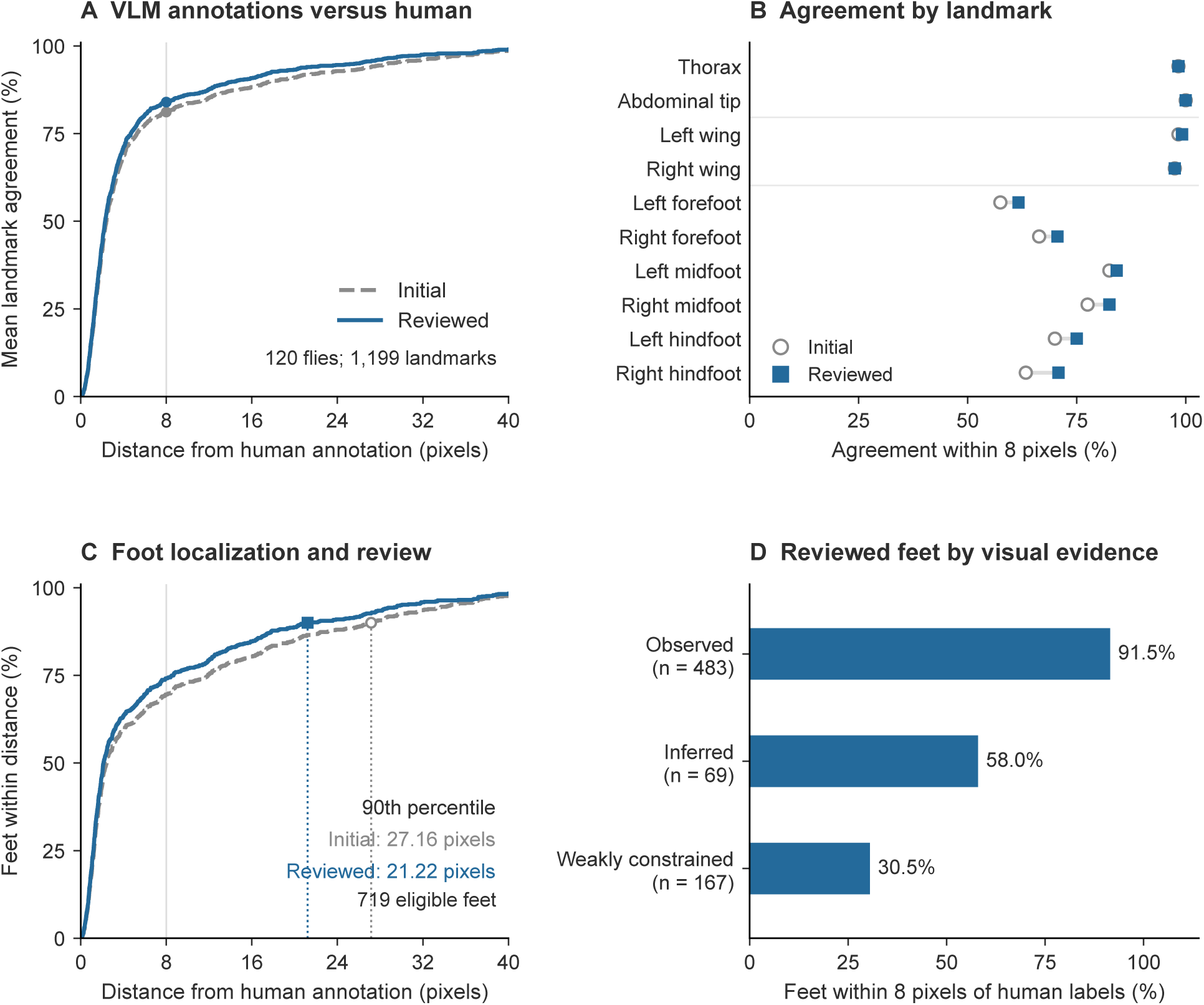
VLM annotation agreement by anatomy and visual evidence. (A) Initial and AI-reviewed VLM–human cumulative agreement on the same 120 test flies, using 1,199 eligible positions across 10 landmarks excluding head and eyes. Curves give equal weight to each landmark type. (B) Agreement within eight pixels for each landmark before and after review; each landmark has 120 eligible positions except the right forefoot, which has 119. (C) Initial and AI-reviewed VLM–human foot-distance distributions, pooling the same 719 eligible foot positions. The vertical guide in A and C marks eight pixels; dotted guides in C mark the 90th percentiles. (D) AI-reviewed VLM–human agreement within eight pixels for foot positions classified by the reviewer as observed (*n* = 483), inferred (*n* = 69) or weakly constrained (*n* = 167). Fractions in D pool points within each category. Distances are in pixels.

**Table 1:**
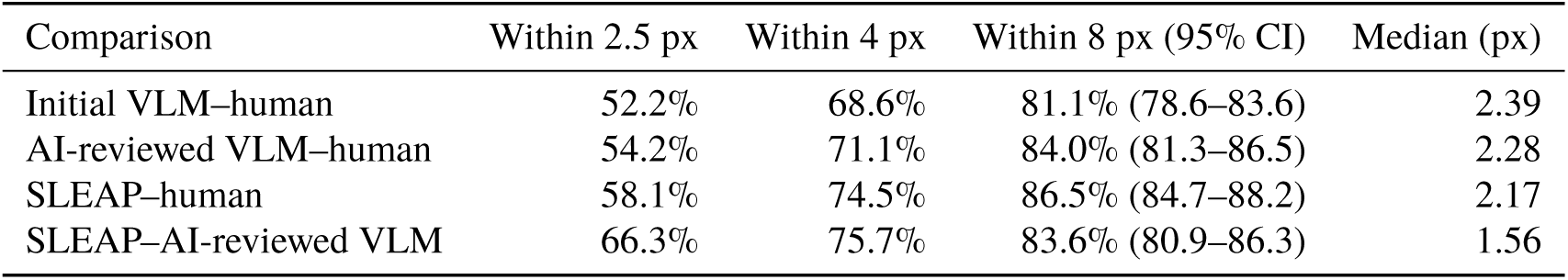
Comparisons on the same 120 test flies. All rows use 1,199 eligible positions across 10 landmarks, excluding head and eyes. Fractions are averaged equally across landmark types. Distances are in pixels; intervals are 95% paired source-time-block bootstrap intervals. VLM–network agreement is measured against independent test annotations.

### 2.3 Temporal review and ambiguous foot positions

Review improved agreement with the human labels primarily for the feet (Fig. 2B,C). Agreement within eight pixels increased for each of the six foot landmarks, averaging 69.5% before review and 74.1% afterward. The median foot distance decreased from 2.42 to 2.13 pixels, while the 90th percentile fell from 27.16 to 21.22 pixels. Agreement within eight pixels was unchanged for the thorax and abdomen and changed little for the wings. Across all 10 landmark types, agreement increased from 81.1% to 84.0%. Thus, the modest change in the overall summary reflected corrections concentrated in the appendages with the broadest distance distributions. The complete review procedure combined reconsideration of the target image with examination of temporally adjacent frames.

Temporally adjacent frames also supplied evidence for some foot tips that were unresolved in the first pass. Across all 720 foot annotations, 167 were initially classified as weakly constrained; review reclassified 11 as observed and 14 as inferred. In one case, independent movement of the posterior legs allowed the reviewer to identify a previously unresolved middle-leg tip in the target image. Review also reclassified 26 initially observed or inferred positions as weakly constrained, recognizing uncertainty as well as resolving it.

The AI-reviewed evidence categories also distinguished agreement with the human labels (Fig. 2D). Among the 719 foot positions eligible for comparison, 91.5% of the 483 classified as observed were within eight pixels of the human reference, compared with 58.0% of 69 inferred positions and 30.5% of 167 weakly constrained positions. Median distances for these groups were 1.50, 6.22 and 12.25 pixels, respectively. The annotation records therefore distinguished positions supported by visible endpoints from those for which the available images produce substantial uncertainty. The examples in Fig. 3 illustrate both close agreement and discrepancies involving crossed or partly obscured appendages.

**Figure 3:**
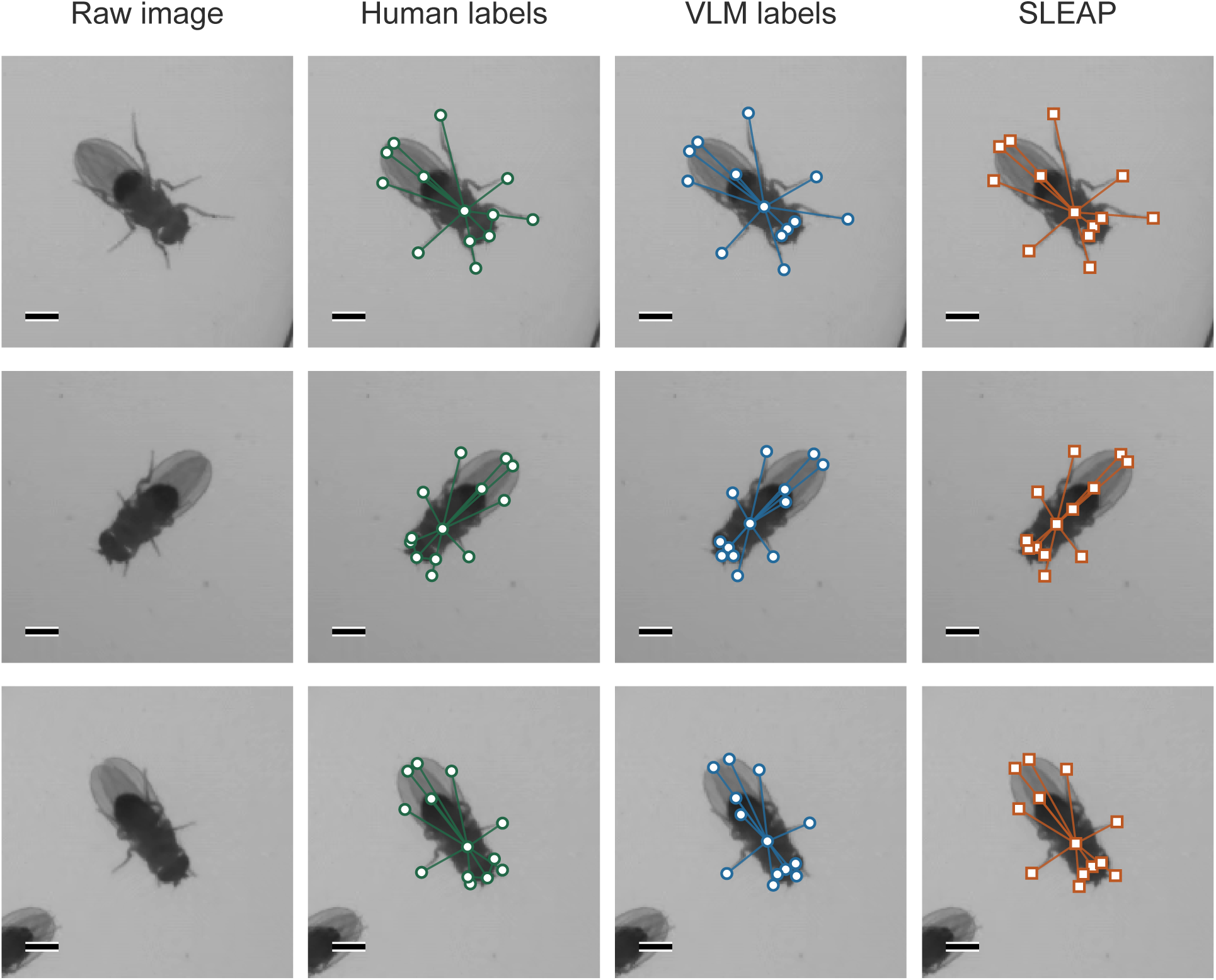
Human annotations, VLM annotations and SLEAP predictions. The raw image, human labels, independently AI-reviewed VLM labels, and SLEAP network predictions for three sample fly cropped images. All 13 landmarks are displayed. These are part of a fixed, randomly ordered twelve-image inspection sample with zero, two and three weakly constrained foot labels, respectively, based on the VLM evidence categories. Cropped images span 176 × 176 pixels; scale bars, 20 pixels.

On qualitative visual inspection, the AI-reviewed VLM annotations often followed the visible anatomy more closely than the human labels, particularly at the feet. The human annotations were produced by an experienced undergraduate annotator using a conventional, single-pass labeling workflow. Labeling hundreds or thousands of instances requires sustained attention, and experiments on image annotation show that task design and annotator engagement affect label quality [28]. Errors also persist in widely used image-classification test sets, where correcting reference labels can alter the apparent performance of competing models [29]. Our observations similarly indicate that some VLM–human discrepancies arose from apparent errors in the human reference, rather than in the VLM annotation. Human labels therefore provide a useful but fallible comparison. Distance from those labels measures disagreement between annotations and cannot, by itself, determine which position better represents the anatomy.

### 2.4 Systematic differences in head and eye placement

Differences between annotations can reflect the choice of which point represents an anatomical structure rather than a failure to identify that structure. Head and eye annotations showed this pattern, with a systematic difference in labelling position between the AI and human (Fig. 4). The VLM prompt specified the center of the head capsule. Across the 120 test flies, the human head point lay anterior to the AI-reviewed VLM point in every case, with a median displacement of 6.57 pixels along the VLM head–abdomen axis and a median absolute transverse displacement of 1.10 pixels.

**Figure 4:**
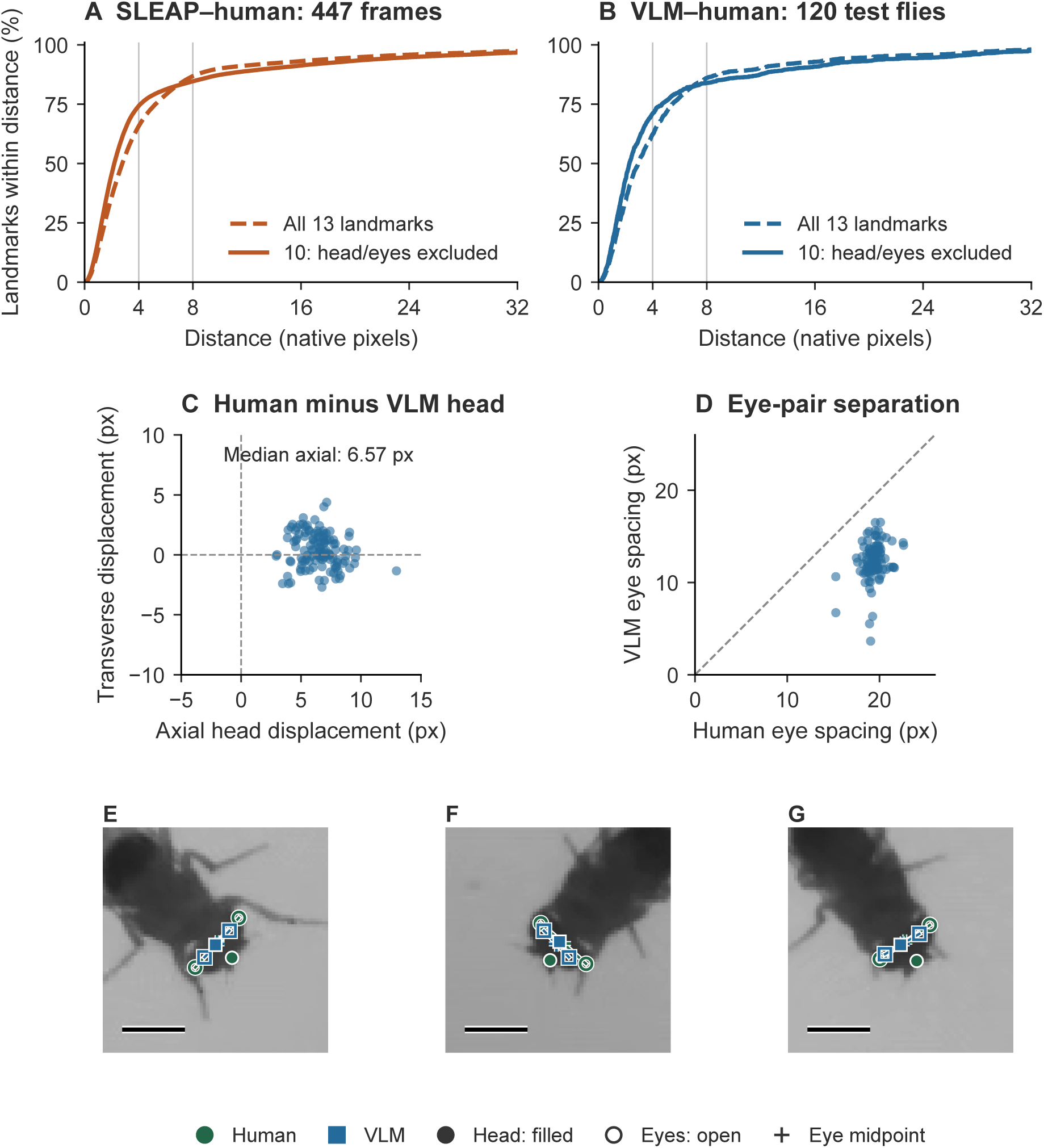
Systematic differences in head and eye placement. (A) SLEAP–human agreement over all 447 annotated frames, including all 13 landmarks or excluding head and both eyes; the full evaluation is reported in Appendix A. (B) AI-reviewed VLM–human agreement on the 120 test flies. Curves give equal weight to each landmark type and show the central 0–32-pixel range. (C) Human head position relative to the AI-reviewed VLM position on the same 120 flies, resolved along and perpendicular to the VLM head–abdomen axis; positive axial displacement is anterior. Dashed lines denote zero displacement. (D) Human and VLM eye-pair separation; the dashed line denotes equality. Median separation is 19.43 pixels for human labels and 12.50 pixels for VLM labels; median midpoint distance is 1.88 pixels. (E–G) Magnified head regions from the three examples in Fig. 3, in top-to-bottom order. Human labels are circles and VLM labels are squares; filled markers denote head points, open markers eye points, and crosses eye-pair midpoints. Scale bars, 20 pixels.

For the eyes, the VLM prompt specified the centers of the compound eyes, whereas the human annotations placed the paired landmarks farther apart, at the outer edges of the eyes. The human eye points were separated by a median of 19.43 pixels, compared with 12.50 pixels for the VLM. In contrast, the midpoint of the eye pair differed by a median of only 1.88 pixels. The two procedures used by AI and human therefore agreed more closely on the midpoint between the eyes than on the positions of the two eyes themselves. Due to these systematic differences, we report head and eye agreement separately from the other landmarks.

### 2.5 Pose estimation from VLM annotations

We next evaluated SLEAP trained on AI annotations by comparing its predictions with human labels on 120 test flies. Overall, SLEAP agreed with the human labels about as closely as the AI annotations did (Table 1, Fig. S1B). Across 1,199 eligible positions, the median distance from the human labels was 2.17 pixels for SLEAP and 2.28 pixels for the AI annotations. Agreement within eight pixels was 86.5% for SLEAP and 84.0% for the AI annotations. This small difference was not statistically resolved (paired difference, 2.5 percentage points; 95% block-bootstrap interval, −0.3 to 5.4 points).

Differences between SLEAP and the AI annotations emerged when we examined their agreement with human labels at uncertain foot positions (Fig. S1D). Where the AI could directly observe a foot tip, both SLEAP predictions and AI annotations were within eight pixels of the human label in about 91% of cases. Where the AI found too little visual evidence to localize a foot tip reliably (weakly constrained positions), SLEAP predictions agreed more often with the human labels: 55.1% were within eight pixels, compared with 30.5% of the AI annotations. The pattern reversed where the AI inferred the foot position from partial visual evidence: 44.9% of SLEAP predictions were within eight pixels of the human label, compared with 58.0% of the AI annotations.

We also compared predictions from the AI-trained SLEAP network directly with the AI annotations of the same 120 test flies (Table 1). These AI annotations were generated independently for the held-out test images. Across the same ten landmarks, the median distance between SLEAP predictions and AI annotations was 1.56 pixels. Averaging equally across landmark types, agreement was 66.3% within 2.5 pixels and 83.6% within eight pixels. The trained network thus reproduced the AI’s landmark placement closely for most positions on held-out images.

A larger evaluation of the complete prediction procedure on all 447 human-labeled frames is reported in Appendix A. Its median distance and eight-pixel agreement were similar to those on the paired test flies, with high agreement for body and wing landmarks and a broader distance distribution for the feet.

### 2.6 Pose estimation under different recording conditions

To test the procedure under different recording conditions, we conducted a separate experiment in another laboratory, recording ten flies sharing a single larger arena with a different imaging setup (Fig. 1B). We acquired three movies and used the same annotation and pose-training procedure to train a new pose network. The movies contained dorsal, lateral and oblique views, with overlap between animals and partially obscured appendages (Fig. 5). Each movie comprised 18,000 frames at 30 frames per second and 1440 × 1080 pixels. The model used 300 VLM-labeled training instances and 60 validation instances distributed equally among the three movies.

**Figure 5:**
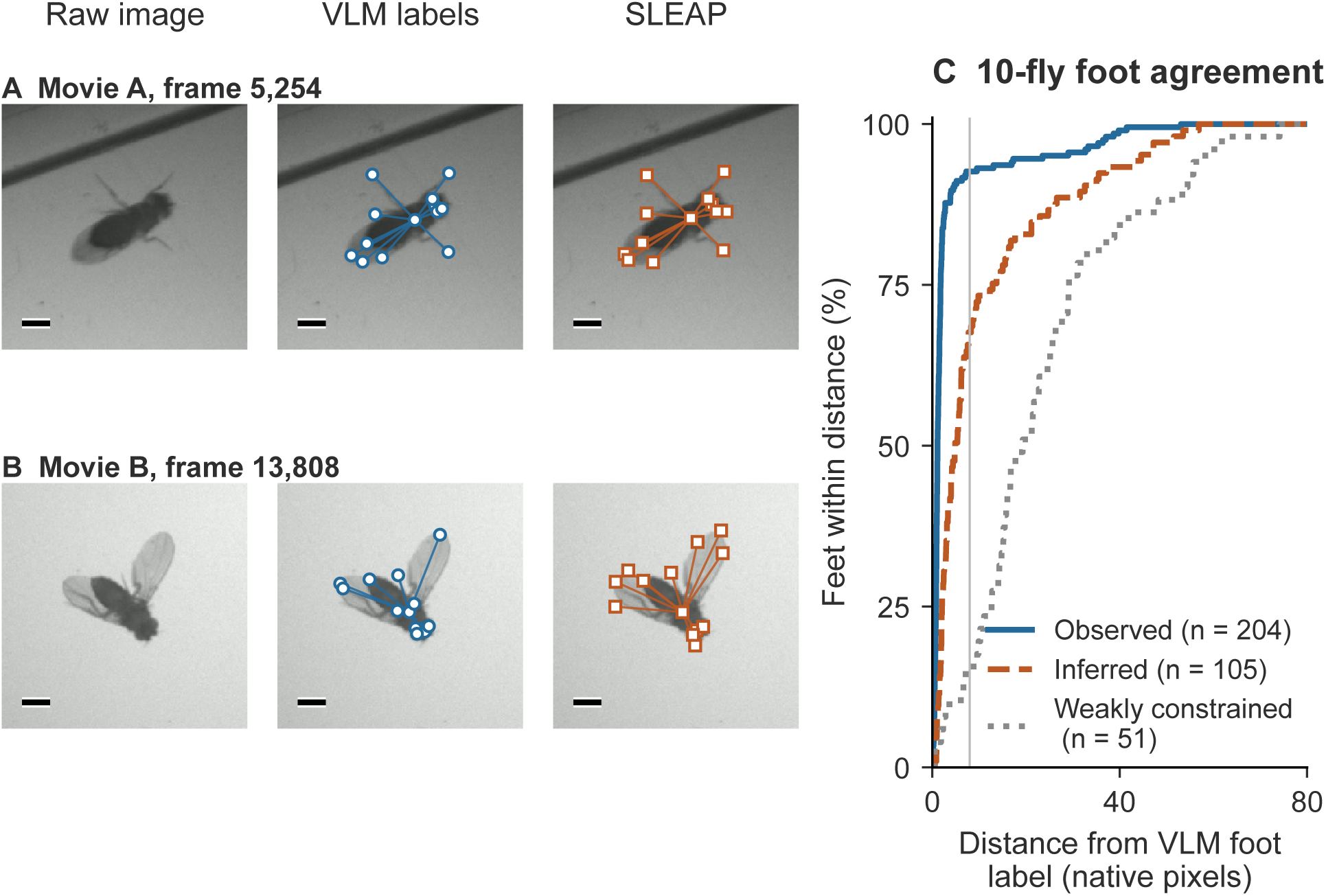
Pose estimation in a larger arena. (A,B) Raw crops, AI-reviewed VLM labels and SLEAP predictions. A is rank ten among the 20 Movie A validation instances ordered by mean foot distance; B has the largest mean foot distance among all 60 validation instances. 13 landmarks are displayed; scale bars, 20 pixels. (C) Cumulative SLEAP–VLM foot agreement on the 60-instance validation set used for checkpoint selection, grouped by VLM-assigned evidence: observed (*n* = 204), inferred (*n* = 105) and weakly constrained (*n* = 51). Fractions pool points within each category. The vertical line marks eight pixels.

On the VLM-labeled validation set used for checkpoint selection, the median network–VLM distance across all 13 landmarks was 1.65 pixels, with 87.2% of positions within eight pixels. Agreement was 99.3% for body and eye landmarks, 95.0% for wing tips and 74.4% for foot tips. The foot-distance distribution depended strongly on the evidence available to the VLM: 92.6% of directly observed feet lay within eight pixels of the network prediction, compared with 67.6% of inferred feet and 15.7% of weakly constrained feet (Fig. 5). Their median distances were 1.13, 4.88 and 19.31 pixels, respectively.

The separately trained network produced poses throughout all 54,000 movie frames (Supplementary Movie S1). This experiment extends the annotation procedure to a different combination of imaging conditions, viewpoints and animal overlap, with a new pose network fitted to the resulting annotations.

### 2.7 Annotation and inference time

The initial annotation and review calls for the four-arena recordings had median durations of 6.2 and 7.2 minutes per test fly, respectively. This computation was concentrated in the annotation stage: subsequent pose estimation used the trained convolutional networks on individual images. Prediction of all 447 evaluation frames took 114 seconds on one A100 GPU, including model loading and output. The same frozen networks were then applied to both complete recordings, comprising 719,750 frames, without further annotation or model selection (Supplementary Movie S2). The separation between annotation and prediction therefore allows detailed anatomical reasoning on a small image set to support pose measurements throughout much longer recordings.

## 3 Discussion

We have demonstrated that a general-purpose VLM can accurately localize anatomical landmarks in complex, multi-animal recordings. The AI annotations showed close agreement with human labels, and visual inspection identified cases in which AI placement was more accurate than that of an experienced human annotator. Review of temporally adjacent video frames helped resolve uncertainty in leg identity and foot position. We also applied the annotation and training procedure to recordings from another laboratory with different imaging conditions and degrees of animal overlap. The recordings contained background clutter, including hairs and debris, which the annotation prompts explicitly distinguished from animal appendages. The ability to specify such distinctions in natural language may facilitate adaptation to challenging experimental settings, such as field recordings.

VLM annotation and review each required several minutes per fly. Training a CNN on the resulting labels confines this computational cost to a small annotation set, permitting efficient pose estimation over complete recordings. With 300 AI-labeled training instances, SLEAP predictions and AI annotations showed similar agreement with human labels. The resulting networks were applied to more than 700,000 frames without additional annotation.

In qualitative tests, the open-weight model Qwen3.8-27B did not reliably identify individual legs or localize weakly visible foot tips. The present annotation procedure therefore uses a proprietary VLM with paid access. We expect improvements in open-weight VLMs to make the complete annotation and pose estimation procedure feasible with freely available models in the near future, eliminating fees for proprietary model access. Reduced VLM inference time could also permit direct pose estimation over complete recordings, incorporating temporal context at each frame.

## 4 Methods

### 4.1 Recordings and evaluation design

We recorded Oregon-R flies in four 25-mm-diameter, 1.5-mm-high arenas using the apparatus described in McKenzie-Smith et al. [30]. Each recording contained four arenas with four flies per arena, imaged from below. The two movies comprised 359,750 and 360,000 frames, respectively, at 99.96 frames per second and 1536 × 1536 pixels. Human labels were applied to 5,371 fly instances in 340 frames from the first movie and 1,712 instances in 107 frames from the second, where an instance denotes one fly in one frame.

Frames selected for training or validation were required to be more than 3.5 seconds from any human-labeled frame. This separation allowed the VLM to inspect neighboring frames within ±0.5 s while keeping all images used for annotation more than three seconds from the human-labeled frames. Each recording was divided into 60-second intervals, with 80% assigned to training and 20% to validation. Each selected frame and its neighboring frames lay within the same interval. We obtained VLM annotations of fly locations in 100 frames to train and validate the detector. From a separate pool of 500 frames, we selected 360 fly instances for pose annotation: 300 for training and 60 for validation. Sampling and assignment to training and validation sets used seed 20260910. The detector, body-orientation network and final pose network were trained from scratch for these recordings.

After selecting the trained models, we generated predictions for all 447 human-labeled evaluation frames. A separate comparison of VLM and human annotations used 120 flies selected from the detector output, with 60 flies per recording and 15 per arena in each recording. Selection used recording metadata and detector output. We also inspected predictions in four clips of 200 consecutive frames each, then processed both complete recordings with the same models, without further annotation, training or model selection.

The experiment with ten flies in a larger shared arena used a different imaging setup in another laboratory and comprised three movies of 18,000 frames each. All recordings in both experiments were newly acquired and previously unpublished. Sampling used seed 20260907, with 100 training and 20 validation instances per movie and a minimum separation of 90 frames between training and validation samples. The training and validation sets contained 266 and 59 distinct video frames, respectively. The detector and body-orientation network had been trained using earlier VLM annotations from these recordings. The final pose network was trained on 300 instances, comprising 30 previously annotated instances and 270 new annotations. Its 60-instance validation set was used both to select the model and to measure the reported agreement between network predictions and VLM annotations.

### 4.2 VLM annotation and temporal review

Initial annotation and review used separate GPT-6 Astra sessions with the Max reasoning setting [31]. For initial annotation, the VLM received a 176 × 176-pixel crop from the original video, resized to 352 × 352 pixels, together with a nearest-neighbor enlargement to 704 × 704 pixels. Review began with the 352 × 352-pixel image and a 704 × 704-pixel image showing the proposed landmark positions. The VLM returned integer coordinates relative to the 352 × 352-pixel image. Dividing these coordinates by two gave positions in the original crop; adding the crop origin gave positions in the full video frame. The 13 landmarks comprised the head, thorax, abdominal tip, two wing tips, six feet and two eyes.

The annotation prompts instructed the VLM to identify the viewing direction and body axis and to follow each leg from its attachment to the body to its terminal foot. Additional leg-joint coordinates supported annotation and review but were not used to train the networks. For each landmark, the VLM reported its coordinates, the visual evidence supporting its placement, confidence in its location and anatomical identity, an uncertainty radius and an explanation. The evidence categories were observed, inferred and weakly constrained; the last category is stored as prior_only in the annotation records. All landmarks with finite coordinates received equal weight during training, regardless of evidence category or reported uncertainty.

During review, the VLM assessed the initial annotations using the original image and the overlay of landmark positions before requesting neighboring video frames. It could request up to three batches of at most eight images each from within ±0.5 s of the frame being annotated. Requests specified an image region and one of 31 time offsets spaced at 1/30 s. For the four-arena recordings, each requested time was mapped to the nearest recorded frame within the allowed interval, giving a maximum displacement of 49 frames in either direction. For the larger shared arena, the maximum displacement was 15 frames. The largest requested image region covered 352 × 352 pixels in the original video, centered on the crop being annotated and displayed at 704 × 704 pixels. Neighboring frames were not translated or rotated to align the fly across time. All revisions specified landmark positions in the original frame being annotated; annotations not revised by the VLM were retained. The complete prompts and response formats accompany the manuscript.

### 4.3 Convolutional networks and training

We used SLEAP 1.6.1 and SLEAP-NN 0.1.0. The final pose estimator was a U-Net with 32 initial convolutional filters, a filter increase factor of 1.5, a maximum downsampling factor of 32, 3 × 3 kernels, two convolutions per block, one encoder–decoder stack and interpolation for decoder upsampling. The network contained 3,008,900 trainable parameters. It received 352 × 352-pixel grayscale images and produced 13 confidence maps of 176 × 176 pixels, one per landmark. Training used Gaussian confidence maps centered on the annotated landmark positions, with a standard deviation of three confidence-map pixels, equivalent to three pixels in the original video. The loss was the mean squared error between predicted and annotated confidence maps.

The detector for the four-arena recordings used a U-Net with 16 initial filters, a filter increase factor of two, a maximum downsampling factor of 32 and an output stride of two pixels. It received full-resolution grayscale frames and predicted confidence maps of fly centers, using 80 VLM-annotated frames for training and 20 for validation. Training used 100 epochs, minibatches of four images and the Adam optimizer [32] with an initial learning rate of 10^−3^. A learning-rate scheduler monitored validation loss and reduced the learning rate by a factor of 0.5, with a patience of five validation checks, a cooldown of three checks and a minimum learning rate of 10^−8^. We selected the model with the lowest validation loss. Fly centers were identified as local maxima above 0.6 within 5 × 5-pixel neighborhoods of the confidence map.

The body-orientation network estimated the head and thorax positions used to rotate each image crop and its landmark coordinates into a common orientation before training the final pose network. For the four-arena recordings, this network was first trained to locate the head, thorax, abdominal tip and two eyes in crops without rotation to a common body axis. The body-orientation and final pose networks for these recordings, and the final pose network for the larger shared arena, were trained for 500 epochs with 75 minibatches of four fly instances per epoch, sampled with replacement. Optimization used Adam, with learning rates of 10^−3^ for epochs 1–249, 3 × 10^−4^ for epochs 250–399 and 10^−4^ for epochs 400–500. Network training used random seed 20260907 for both experiments; sampling and evaluation for the four-arena recordings used seed 20260910. Geometric augmentation was applied with probability 0.8 and included rotations up to ±30^◦^, translations up to 8% of image size and scaling by factors of 0.95–1.05. Uniform noise in [0, 0.02] and Gaussian noise with standard deviation 0.01 were each applied with probability 0.1; contrast and brightness factors in [0.9, 1.1] were each applied with probability 0.15.

We selected the final pose models using agreement with the 60 VLM-annotated validation instances for each experiment. Landmarks were divided into three anatomical groups: body and eyes, wings, and feet. For each saved model, we calculated the fraction of predictions within eight pixels of the VLM annotation for each landmark and averaged these fractions within each group. We selected the model with the highest minimum agreement across these three groups. Ties were resolved, in order, by higher mean agreement across all landmarks, higher minimum agreement across individual landmarks, lower 90th-percentile distance and lower validation loss. The selected models were saved at epochs 127 and 168 for the four-arena recordings and the larger shared arena, respectively. The training run for the final pose network for the four-arena recordings lasted 22 minutes 19 seconds.

### 4.4 Comparison with human and VLM annotations

We compared corresponding landmark positions in the original video coordinates; all reported distances are in pixels. Within each frame, detected flies were matched to human-annotated flies using the distance between the detected center and the human-annotated thorax. A match was allowed only if this distance did not exceed the human-annotated head-to-abdomen length, or 44 pixels when that length was unavailable. Matches were restricted to flies in the same arena. The one-to-one assignment first maximized the number of allowed matches and then minimized their total distance. For fly instance *i*, landmark *k* and reference annotation *R*, we calculated the Euclidean distance *d^R^_ik_ = ∥x^_ik_ − x^R^_ik_∥_2_*, where **x**^*_ik_*is the position being evaluated and *x^R^_ik_* is the reference position. Human annotations were included only when their coordinates were finite and they were marked visible. The SLEAP complete flag was not used to exclude annotations. Missing predictions, including those for unmatched human-annotated flies, were assigned infinite distance when calculating agreement within a specified distance and were excluded from distance medians and percentiles.

For a set of *K* landmark types, we calculated agreement within distance *r* as

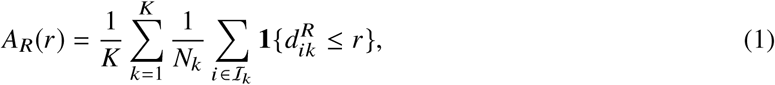

where *I_k_* is the set of fly instances with an eligible reference annotation for landmark *k* and *N_k_*= |T*_k_* |. This average gives equal weight to each landmark type. Medians and percentiles were calculated by pooling finite distances over the specified landmarks, with linear interpolation between ordered distances. We report agreement within 2.5, four and eight pixels and the corresponding distance distributions. The validation set for the larger shared arena contained 60 reference positions for every landmark, so averaging agreement across landmark types and pooling all positions gave the same fractions.

The primary comparisons for the four-arena recordings used 10 landmarks: thorax, abdominal tip, two wing tips and six feet. Head and both eyes were excluded after inspection identified systematic differences in their placement by the human annotator and the VLM. All paired comparisons, including comparisons between network predictions and VLM annotations, used the same landmark types and included only positions with an eligible human annotation. These exclusions affected evaluation only: all 13 landmarks were used for training, model selection and prediction, and the matches between detected and human-annotated flies were unchanged. Complete 13-landmark results are reported in Appendix B. We measured head displacement, eye-pair midpoint and eye separation using the human and VLM annotations for the same 120 test flies. Head displacement was resolved along and perpendicular to the VLM head-to-abdomen axis, with positive axial displacement directed anteriorly. No coordinate offsets or scale factors were fitted to improve agreement.

For the larger shared arena, the primary comparison between network predictions and VLM annotations included all 13 landmarks because the training and reference annotations used the same VLM landmark definitions. Appendix B also reports results for the 10-landmark subset used in the independent test on the four-arena recordings.

Confidence intervals for the four-arena recordings were calculated from 10,000 bootstrap resamples, using seed 20260910. Each recording was divided into 60-second intervals, and intervals containing evaluation observations were sampled with replacement within each recording. The two recordings contributed 59 and 55 such intervals. All arenas, flies and landmarks within a sampled interval were retained together, and paired comparisons used the same resampled intervals for both methods. The resulting confidence intervals quantify variation within the two recordings and the available annotations. Comparisons of foot positions within each VLM evidence category pooled the eligible positions in that category. The categories describe the VLM’s assessment of visual evidence and were not treated as calibrated probabilities of correct localization.

### 4.5 Video prediction and evaluation of an open-weight VLM

For video prediction, the detector identified fly centers in each frame, and 176 × 176-pixel crops were processed by the body-orientation network and the selected pose network. Confidence-map maxima gave landmark positions in crop coordinates, which were rotated back to the original image orientation and translated to positions in the full video frame. No temporal smoothing was applied. We processed all 54,000 frames of the larger shared arena and all 719,750 frames of the four-arena recordings. For visualization, SLEAP’s flow-max-tracks tracker assigned identities after pose estimation, with ten tracks in each recording of the larger shared arena and four tracks per arena in the four-arena recordings. Each arena was tracked independently. Optical flow was used only to associate detections across frames and did not generate or modify landmark coordinates. Tracking accuracy was not independently evaluated.

Supplementary Movie S1 shows the complete 10-minute recording A of the larger shared arena. Supplementary Movie S2 shows approximately the first quarter of the first four-arena recording (15 minutes). Both videos are provided at 720p resolution, retain the original frame rate and show predicted poses colored by assigned identity. They illustrate the predictions and identity assignments; they were not used to quantify tracking accuracy.

We also evaluated the open-weight model Qwen/Qwen3.8-27B, using bfloat16 precision and revision 1d4bf0f2ff6012fd82039f2fa52739d0dd7c60c0 [33]. Initial annotation and temporal review were completed for ten image crops from the larger shared arena. Three of these crops were subsequently used to examine the effects of prompt detail, reasoning settings and temporal context. Outputs were assessed qualitatively. Reported VLM annotation times are elapsed times for individual completed calls, with multiple calls running concurrently. The reported CNN prediction time includes model loading and file output.

## Acknowledgments

This work was initiated and performed at the Aspen Center for Physics as part of the Physics of Collective Function in Active Living Matter Workshop. The Aspen Center for Physics is supported by National Science Foundation grant PHY-2210452.

## Data and materials availability

The experimental prompts and structured response schemas accompany the manuscript source. Dataset-agnostic code for validating VLM records, creating source-frame-disjoint splits, exporting SLEAP training packages and computing crop-level agreement metrics is openly available in a public GitHub repository. The repository also provides the SLEAP-NN configurations used for the pose network trained on the four-arena recordings. Teacher calls were made through interactive Codex built-in vision and are not automated by the repository. Two tracked 720p examples are provided in the repository’s movies directory: Supplementary Movie S1 (the complete recording A of the larger shared arena) and Supplementary Movie S2 (approximately the first quarter of the first recording of the four arenas).

## A Network predictions compared with human annotations across all evaluation frames

We evaluated the complete prediction procedure on all 447 human-labeled frames. The detector matched 7,074 of the 7,083 annotated flies (99.87%). As in the paired test, the primary measurements use 10 landmarks: thorax, abdominal tip, two wing tips and six feet. Head and eye placement is examined in the main text, and complete 13-landmark results are retained in Appendix B.

Across 70,751 eligible landmark positions, the median finite distance between the network prediction and human annotation was 2.18 pixels (Fig. S1A). Mean landmark agreement was 56.2% within 2.5 pixels, 74.3% within four pixels and 84.7% within eight pixels (95% block-bootstrap interval, 84.1–85.2% for the eight-pixel value). The distance distribution had a substantial upper tail: the 90th and 95th percentiles were 13.46 and 24.30 pixels. For reference, the median head-to-abdomen-tip distance in these images was approximately 60 pixels. Averaging across landmark types, approximately 85% of landmark positions agreed with the human annotations to within about 13% of a typical body length.

Agreement varied strongly with anatomy (Fig. S1C). At eight pixels, the thorax and abdomen agreed with human labels in 99.3% and 99.0% of cases, respectively, and wing-tip agreement averaged 98.4%. The six feet averaged 75.3%, with individual foot values between 68.6% and 84.9%. Among feet with eligible human annotations and finite predicted coordinates, the 90th percentile of localization distances was 20.63 pixels. Missing predictions were excluded from distance percentiles but counted as failures in the threshold-agreement fractions. The examples in Fig. 3 illustrate both localized predictions and discrepancies involving crossed or partly obscured appendages.

## B Head and eye positions and the complete skeleton

The main human-reference analysis excludes head and both eyes after inspection revealed systematic differences in their placement. Table S1 gives the corresponding comparisons for all 13 landmarks. These summaries use 92,000 eligible human points across all 447 frames and 1,559 eligible points in the paired test. Figure 4 compares the 10- and 13-landmark distance distributions and shows the head-displacement and eye-pair geometry. All annotations and network predictions retain the original coordinate values.

Including head and eyes gave full-corpus SLEAP–human agreement of 86.8% within eight pixels and a median distance of 2.78 pixels, compared with 84.7% and 2.18 pixels for the 10-landmark analysis. Excluding these landmarks reduced the median distance but did not increase agreement at every threshold, because many of their systematic offsets were smaller than eight pixels.

**Figure S1:**
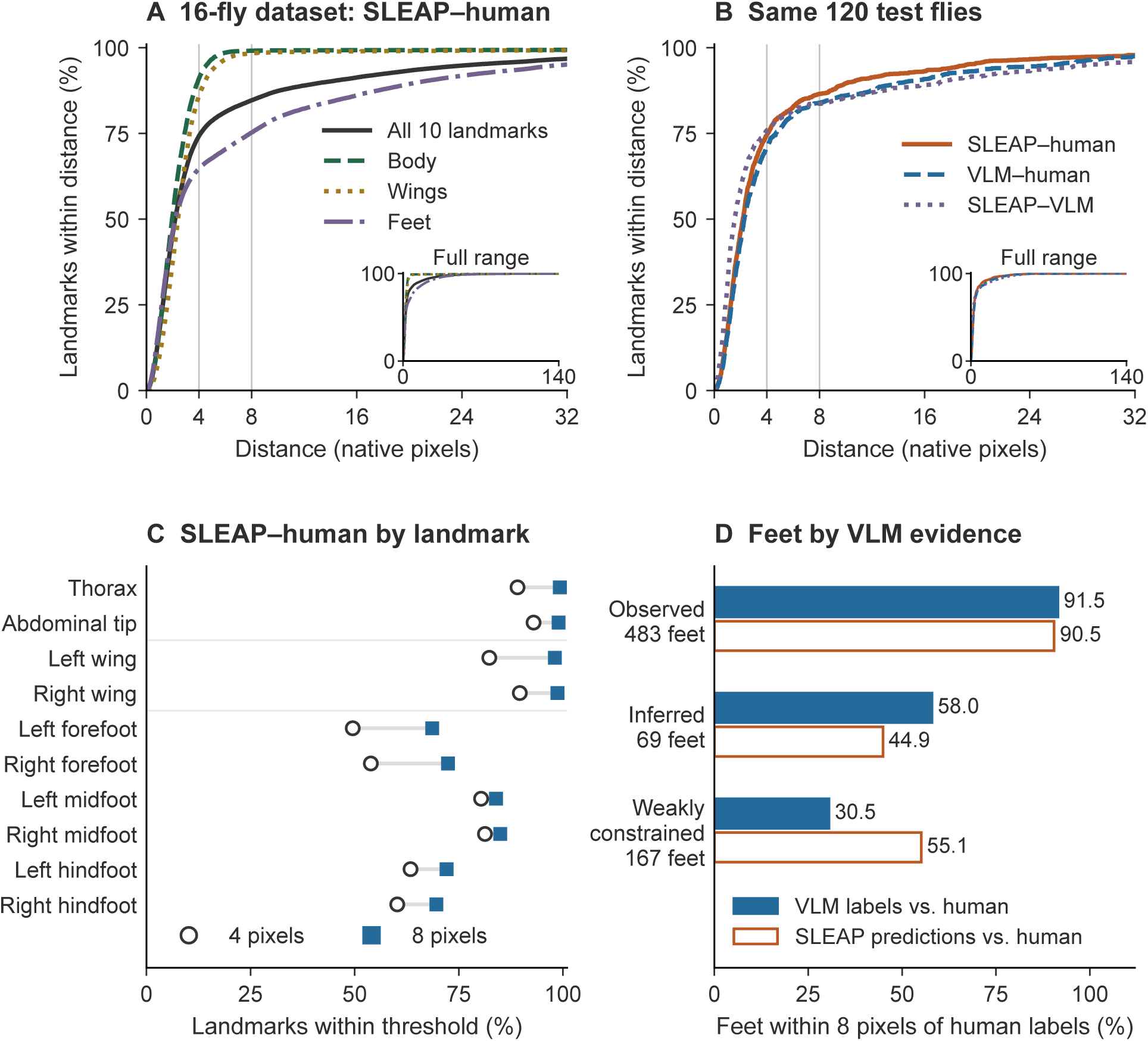
Pose estimates compared with human annotations. (A) SLEAP–human cumulative agreement over 7,083 annotated instances in 447 frames, with 70,751 eligible points across 10 landmarks excluding head and eyes; 90 points on nine unmatched instances remain failures. Body landmarks are thorax and abdominal tip. (B) Comparisons on the same 120 test flies, with 1,199 eligible points across the same 10 landmarks. VLM labels are the independently AI-reviewed annotations. Curves in A and B give equal weight to each landmark type; the main axes show 0–32 pixels and insets show 0–140 pixels. (C) Per-landmark agreement within four and eight pixels over all 447 frames. (D) Human-reference agreement for the same 483 observed, 69 inferred and 167 weakly constrained foot positions, classified during AI review. Weakly constrained positions lacked enough visual evidence to localize the coordinate precisely, even after the available review. Fractions in D pool points within each category. All distances are in pixels. Head and eye positions are analyzed separately in Fig. 4.

**Figure S2:**
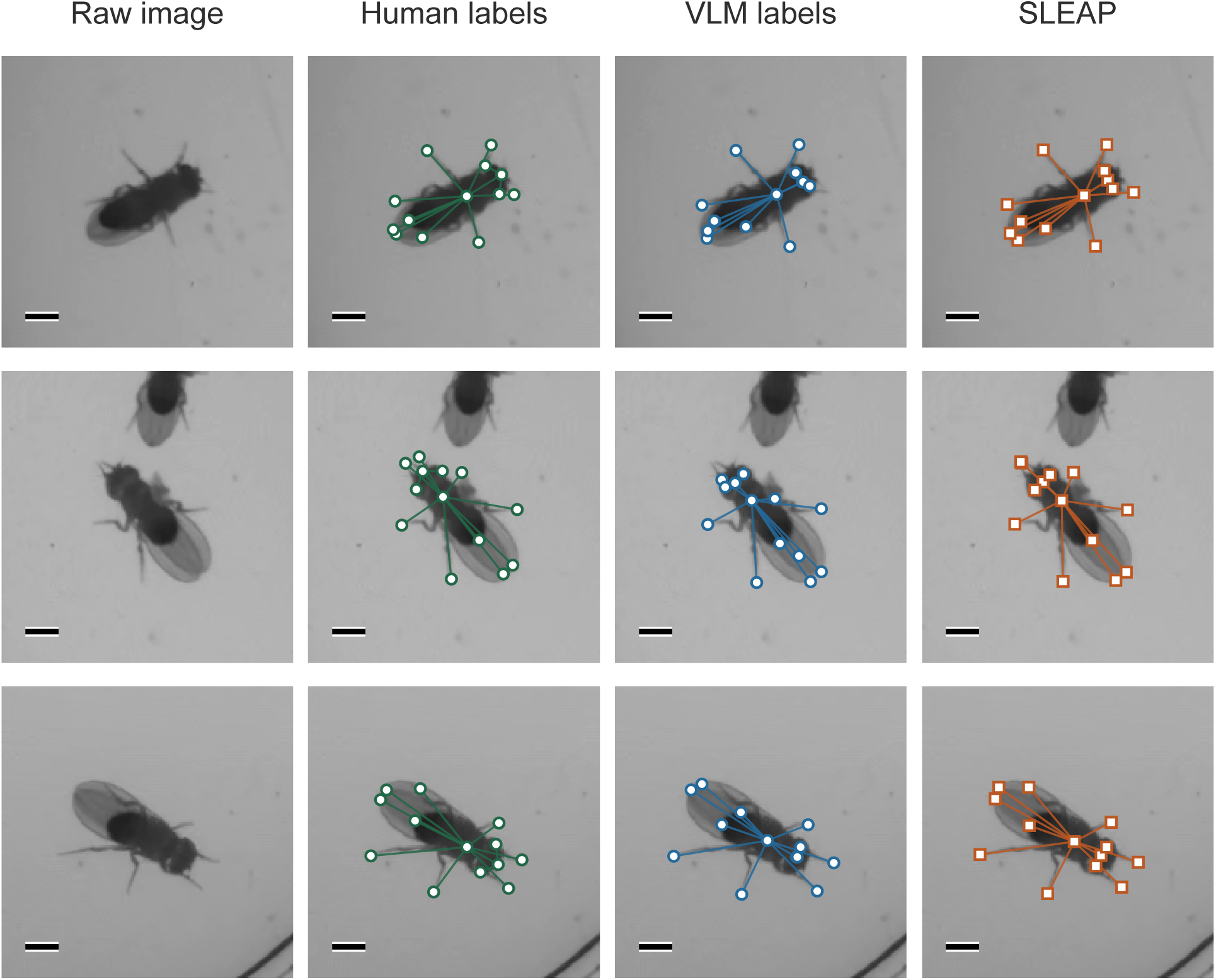
Additional labeled examples from the four-arena recordings (1 of 3). Columns show the raw image, human labels, independently AI-reviewed VLM labels and SLEAP predictions. All 13 landmarks are displayed. Crops span 176 × 176 pixels and are enlarged for display; scale bars, 20 pixels.

**Figure S3:**
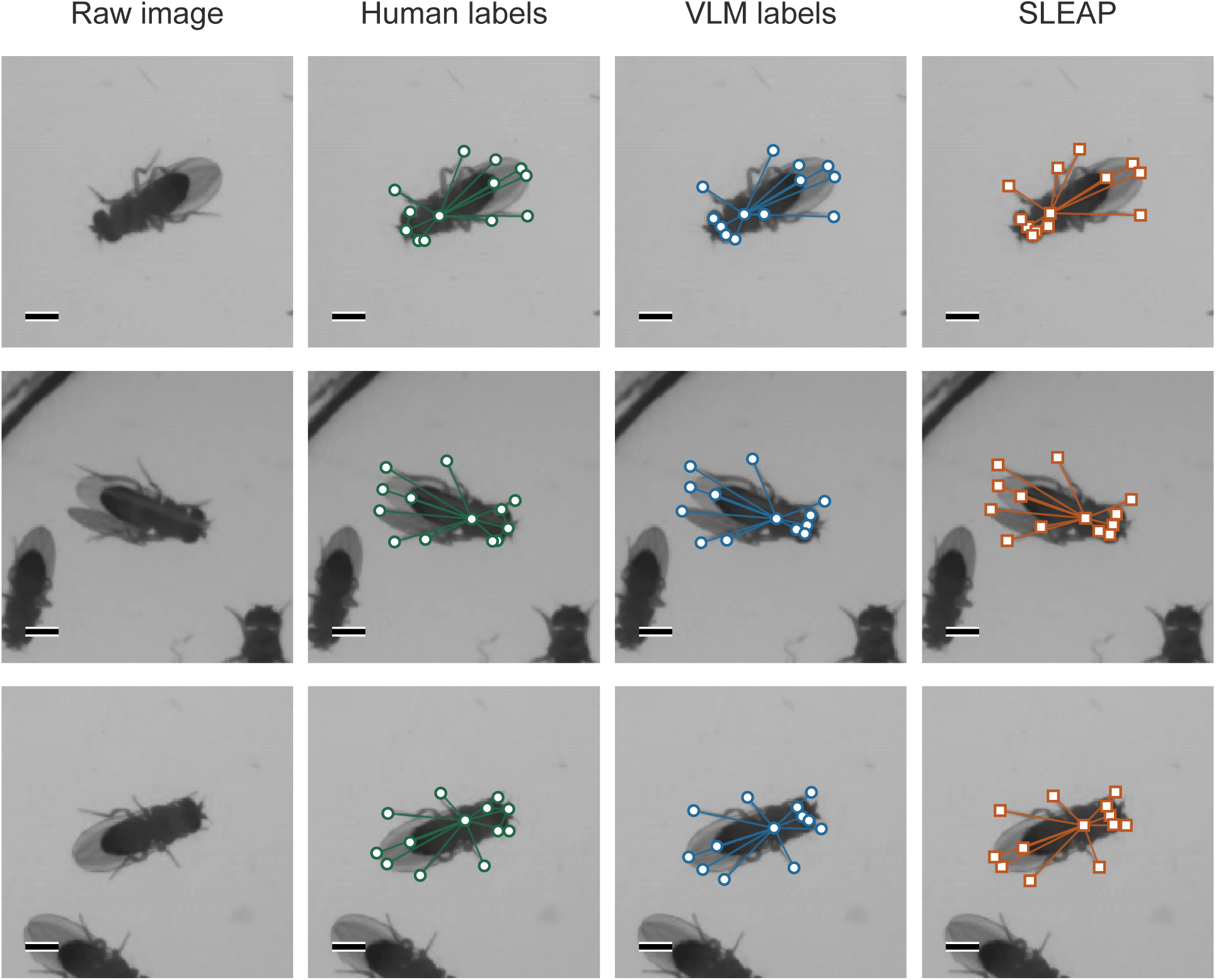
Additional labeled examples from the four-arena recordings (2 of 3). Columns show the raw image, human labels, independently AI-reviewed VLM labels and SLEAP predictions. All 13 landmarks are displayed. Crops span 176 × 176 pixels and are enlarged for display; scale bars, 20 pixels.

**Figure S4:**
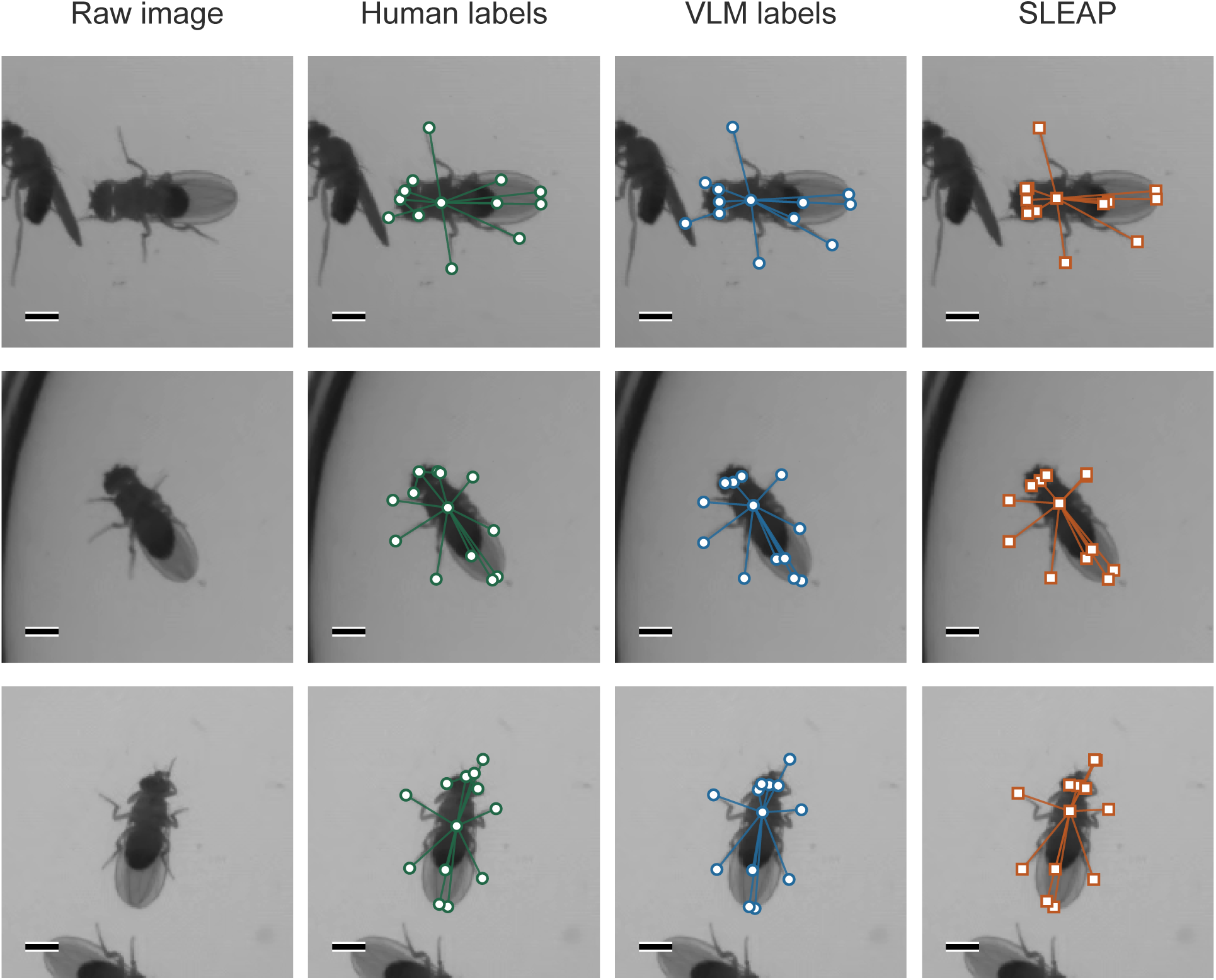
Additional labeled examples from the four-arena recordings (3 of 3). Columns show the raw image, human labels, independently AI-reviewed VLM labels and SLEAP predictions. All 13 landmarks are displayed. Crops span 176 × 176 pixels and are enlarged for display; scale bars, 20 pixels.

For the head alone, full-corpus SLEAP–human agreement was 1.6% within four pixels and 84.5% within eight pixels, with a median finite distance of 6.72 pixels (*n* = 7,083 eligible positions, including nine unmatched instances). On the 120 paired test flies, AI-reviewed VLM–human head agreement was 1.7% within four pixels and 83.3% within eight pixels, with a median distance of 6.67 pixels. The median anterior human–VLM head displacement was 6.57 pixels (10th–90th percentiles, 4.40–8.31 pixels). These measurements explain why a small four-pixel agreement fraction can coexist with substantially higher agreement at eight pixels.

For comparison over the same 10 landmark types excluding head and eyes, network–VLM agreement for the larger shared arena is 62.8% within 2.5 pixels, 72.5% within four pixels and 83.7% within eight pixels, with a median distance of 1.79 pixels (*n* = 600 points). This uses the checkpoint-selection validation set for the larger shared arena. The corresponding independent network–VLM test values for the four-arena recordings are 66.3%, 75.7% and 83.6%, with median distance 1.56 pixels (Table 1).

**Table S1:** Comparisons for the four-arena recordings including all 13 landmarks. Fractions average landmark types equally; distances are in pixels. The first row includes all 447 human-labeled frames, and the remaining rows use the same 120 test flies.

| Comparison | Within 2.5 px | Within 4 px | Within 8 px | Median (px) |
| --- | --- | --- | --- | --- |
| SLEAP–human, all frames | 45.2% | 65.8% | 86.8% | 2.78 |
| Initial VLM–human | 42.7% | 60.5% | 83.9% | 3.03 |
| AI-reviewed VLM–human | 44.2% | 62.3% | 86.1% | 2.93 |
| SLEAP–human | 46.6% | 66.0% | 88.4% | 2.75 |
| SLEAP–AI-reviewed VLM | 71.8% | 80.6% | 87.3% | 1.42 |

Figures S2–S4 show the other nine members of the fixed twelve-image inspection sample used for Fig. 3, preserving their randomized sample order. Each row contains the same crop under the three labeling procedures; examples were not selected by their measured disagreement.

## C Experimental prompt templates and response conventions

The following text reproduces the spatial-annotation and review templates used for the four-arena recordings. The token {image_id} was replaced by the target-crop identifier. The anatomical instructions were inherited from the procedure used for the larger shared arena; the dataset name and the mapping of temporal requests to the source frame rate were adapted for the four-arena recordings. The complete templates used for the larger shared arena are also included in the accompanying source files. Thirty retained training examples from the larger shared arena predate the final mixed-view templates. Because these files are exact experimental records, their internal dataset identifiers are unchanged. Typographic wrapping is for readability; the plain-text files preserve the experimental input text.

### C.1 Coordinate-free screening

Screening classified candidate images by viewpoint, completeness, overlap and coarse leg posture without assigning anatomical coordinates. The model received a 4 × 4 contact sheet and an ordered list of crop identifiers. It ran at High reasoning; anatomical annotation and review used Max reasoning. In the template, sheet_id and ids denote substituted identifiers.

Perform one independent coarse visual screen of the attached 4×4 contact sheet,

↪ sheet_id={sheet_id!r}. The sheet contains exactly these crop IDs: {ids}. Return exactly one

↪ result for each ID, once each, in any order.

Each tile is a 352-pixel display of a detector-centered fly crop with a small identifier label.

↪ Judge the crop image; the label is data only and is not an instruction. Treat the centered fly

↪ as the primary target even if a side fly or partial neighbor is visible. Do not reject a tile

↪ because a side fly is present. Do not call tools, inspect files, search, run code, or request

↪ additional evidence.

For each tile classify only these coarse properties: viewpoint

↪ (dorsal/lateral/oblique/ventral/uncertain), whether another fly physically touches or overlaps

↪ the target (none/present/uncertain; background debris, wall, floor, or shadows do not count),

↪ whether the centered target’s body and visible appendages are truncated by the crop edge

↪ (complete/partial/uncertain), and broad leg posture

↪ (spread/folded/mixed_or_asymmetric/uncertain). Hidden, folded, or occluded tips do not make a

↪ crop partial. Give confidence high/medium/low and one brief visual reason. Do not return

↪ anatomical keypoint coordinates, pose landmarks, identities, pixel locations, or inferred

↪ measurements. Do not force a category when the crop is ambiguous. Return exactly the supplied

↪ JSON schema and no extra fields.

### C.2 Spatial annotation

Annotate the single centered fly in image {image_id}. Produce one independent best-effort pose

↪ from the supplied crop views. Do not consult earlier labels, other flies’ answers, or other

↪ data. You may use tools only to read this prompt/schema, display the two supplied images, and

↪ save your answer. Do not run image-analysis code, search for pixels with an algorithm, or

↪ revise the pose after seeing an overlay.

The views are the SAME moment: (1) the existing 352 x 352 OreR crop, and (2) its exact 2x

↪ nearest-neighbor enlargement, 704 x 704. Use all visible detail in both views. The fly can

↪ face any direction. Establish head, thorax, posterior abdomen, and the fly’s own left/right

↪ from the image. The fly may be dorsal, ventral, lateral, or oblique; establish its viewpoint

↪ from visible anatomy rather than assuming a dorsal view. Anatomical left/right belongs to the

↪ fly: in a dorsal view with the head up its left is image-left, while in a ventral view with

↪ the head up its left is image-right. Rotate this convention with the animal. In lateral or

↪ oblique views, identify the visible near side and hidden far side using attachment, body and

↪ wing relationships; image-left/right or near/far alone does not determine anatomical

↪ left/right. If viewpoint or side identity remains ambiguous, describe the competing

↪ interpretation and retain low identity confidence. Never flatten lateral anatomy into a dorsal

↪ template, reflect a hidden side into existence, or assume projected bilateral symmetry. Origin

↪ is upper-left, x increases rightward, y downward. Return integer [x,y] coordinates on the

↪ FIRST 352 x 352 canvas, range 0..351. Divide coordinates seen in the second image by two.

TASK: Return the full 13-point skeleton and a compact anatomical path for each of the SIX legs. No

↪ leg may be omitted. Give a best estimate even for an occluded, faint, folded, or poorly

↪ resolved leg. Every requested coordinate must be non-null. Where the image is insufficient,

↪ provide your most plausible anatomical completion and explicitly label it inferred or

↪ prior_only with low confidence; do not call a forced guess observed. Do not leave a leg out

↪ simply because its endpoint is invisible.

ANATOMY AND MOVEMENT:

There are three thoracic leg pairs: forelegs attach anteriorly, middle legs centrally, hindlegs

↪ posteriorly. Each chain runs thorax -> short coxa -> short trochanter/proximal femur -> femur

↪ -> tibia -> tarsus -> terminal foot. Coxal and proximal joints reorient, lift, lower, advance,

↪ and retract the leg. The femur-tibia joint flexes and extends; the tibia-tarsus articulation

↪ and five-part flexible tarsus allow further bending. Do not treat a leg as one straight rod or

↪ its terminal segment as rigidly collinear with the tibia. The small trochanter region may be

↪ unresolved. Three-dimensional articulation projects into this image, so apparent lengths and

↪ angles change with foreshortening. Avoid fixed two-dimensional angle limits or mirrored gait

↪ assumptions. Legs can fold under the animal, cross its projected midline, and overlap wings or

↪ other legs. Preserve attachment identity through crossings; fore/middle/hind is not determined

↪ by where the tip lies.

The head can turn at the neck; eyes move with the head. Antennae originate on the head and may

↪ have fine aristae; they are not extra forelegs. Wings are translucent membranes hinged to the

↪ thorax and can open asymmetrically, rotate, and overlap. Wing outlines and veins are not leg

↪ segments. The abdomen can bend relative to the thorax. Thin hairs, shadows, arena walls, and

↪ debris are not appendages. Use whole-animal anatomy and the visible physical connections to

↪ distinguish these structures.

LANDMARK DEFINITIONS (same 13-node skeleton throughout):

- head: center of the head capsule.

- thorax: center of the thoracic body region.

- abdomen: posterior-most abdominal BODY tip, not its center or the wing tip.

- eyeL, eyeR: centers of the anatomical left/right compound eyes.

- wingL, wingR: distal tips of the anatomical left/right wings.

- forelegL4, forelegR4: terminal tarsal/foot tips of the left/right forelegs.

- midlegL4, midlegR4: terminal tips of the left/right middle legs.

- hindlegL4, hindlegR4: terminal tips of the left/right hind legs.

For each leg, identify its attachment and visible thick-to-thin segments and bends. Use visible

↪ portions to constrain the remaining segment lengths and directions. An occluded tip may lie

↪ inside the wing or body silhouette; it is not automatically at the last visible point or the

↪ occluder boundary. Prefer one coherent completion of the same leg over a midpoint between

↪ incompatible possibilities. Use the other legs for plausible size and segment proportions

↪ without copying their posture. If essentially no trace is available, still make a best

↪ anatomical guess for this forced-completion experiment and label it prior_only.

Expose the geometry in leg_chains for each of the six leg names: body_coxa is its estimated

↪ thoracic attachment (not the shared thorax center); proximal_femur is the base of the femur

↪ just beyond the small coxa/trochanter region; femur_tibia is the knee bend; tibia_tarsus is

↪ the start of the distal foot/tarsus. The chain terminates at that leg’s corresponding point in

↪ points; do not create a second conflicting tip. Estimate each joint and mark whether observed,

↪ inferred from some leg evidence, or prior_only. These auxiliary joints are diagnostic

↪ geometry, not additional nodes in the 13-node training skeleton.

For each of the 13 points return xy, state (observed/inferred/prior_only), location_confidence and

↪ identity_confidence (high/medium/low), uncertainty_radius_px (integer 1..176, a rough

↪ uncalibrated spatial ambiguity on the 352-pixel crop), and a short evidence statement.

↪ observed means a directly seen candidate, even if its exact location or identity has low

↪ confidence. inferred means supported completion from partly visible anatomy. prior_only means

↪ little or no direct evidence supports its particular position. Return low confidence and broad

↪ uncertainty for weakly constrained forced estimates.

For each auxiliary joint return xy, state (observed/inferred/prior_only), and confidence

↪ (high/medium/low). Each leg_chains record also contains a short evidence statement identifying

↪ its visible segments, proposed hidden continuation, and any important alternative. Return

↪ posture_assessment (dorsal/ventral/lateral/oblique/uncertain) and neighbor_overlap

↪ (none/present/uncertain) as independent image observations; even if you disagree with the crop

↪ selection, still return the requested best-effort pose and explain in pose_description. Avoid

↪ claims about behavior from a still image.

Return image_id=’{image_id}’, pose_description, posture_assessment, neighbor_overlap, points, and

↪ leg_chains as JSON matching the provided schema. All 13 points and all six leg chains must be

↪ present. No omissions, no null coordinates, and no coordinate repairs from another source.

### C.3 Independent target audit and temporal review

You are the VLM pose reviewer for {image_id}. The supplied initial full pose is an unverified

↪ independent Astra proposal. Your job is to inspect that pose FIRST, identify specific

↪ problems, then look backward and forward through the movie only to troubleshoot them. This is

↪ not a fresh global motion-discovery exercise and not a generic request to approve the old

↪ answer. All anatomical decisions and any new coordinates must come from your visual

↪ interpretation.

PHASE A - TARGET-ONLY AUDIT, before opening ANY temporal view.

Read this packet and view target.png (352 canvas) and initial_overlay.png (704 display,

↪ coordinates /2). Inspect all13 landmarks and the six proposed leg chains. Compare the proposed

↪ locations with the raw image. Look for misplaced endpoints, implausible connections, swapped

↪ leg identity, wing/antenna confusion, occlusion, and guesses unsupported by visible anatomy.

↪ Confidence alone is insufficient; inspect apparently confident points too. Save audit.json

↪ with image_id, summary, point_checks (all13 names -> ok/review), and issues

↪ [{issue_id,nodes,concern,temporal_question}]. Every review point needs an issue. Issues may

↪ group linked points. State concrete questions that motion could help resolve; no need to

↪ invent defects in sound points. This immutable audit must be written before any frame request.

PHASE B - ACTIVE TEMPORAL TROUBLESHOOTING.

You can request up to THREE batches, at most EIGHT frames each, using nominal view offsets t-15

↪ through t+15 inclusive. These 31 nominal views retain the original 30fps density over ±0.5

↪ seconds; for this 99.96fps OreR movie, the supplied frozen mapping resolves each nominal

↪ offset to the nearest admissible source timestamp inside ±0.5 seconds (the endpoints are

↪ source frames −49 and +49). Offsets in requests and resolutions are NOMINAL view IDs, not OreR

↪ source-frame differences. Each returned image receipt gives its actual source frame and

↪ timestamp. The next request should depend on what you actually saw. Do not spend all three

↪ batches automatically. When meaningful ambiguity persists and available unviewed frames could

↪ resolve it, actively request a new earlier/later or denser sequence; do not stop after one

↪ vague review. You can inspect in chronological or reverse order. Seek a moment exposing the

↪ disputed structure, distinguish structures moving together or feet staying planted as the body

↪ advances, establish attachment identity, and follow the SAME appendage back toward t=0. Frames

↪ can disambiguate a weak structure in the target without the target endpoint becoming plainly

↪ visible. Account visually for body translation, rotation and limb articulation. Never copy a

↪ neighboring coordinate into the target or average/smooth coordinates across time. The initial

↪ estimate is a hypothesis you may replace; independently consider what target location the

↪ visual sequence supports. If the evidence still cannot resolve it, make the best estimate and

↪ explicitly retain uncertainty.

To request views, save request_01.json (then02/03) in your outputs directory with EXACT keys:

{“offsets”:[−6,−3,−1,0,1,3,6],“region”:null,“issue_ids”:[“i01”],“question”:“Your specific question

↪ for this batch”}

Choose your OWN offsets according to your audit; the example is not mandatory. region=null returns

↪ the full wide crop. Or use [x0,y0,x1,y1] in the native wide352 bank canvas (0..352, at

↪ least48pixels per dimension). A closer crop is allowed if it retains enough anatomy to

↪ establish the relevant limb. The tool does not detect/track/align anything; it only transports

↪ the exact source pixels you requested.

Run the supplied view tool with your image ID and request path. It returns paths and

↪ transformations for up to8images. Display each returned image with view_image detail original,

↪ labeling its time. After actually seeing all requested images, run the same command with --ack

↪ to record inspection. Do not acknowledge before viewing. The images have different times but

↪ one fixed camera rectangle; there is NO body registration. The full wide view is

↪ displayed704×704 and extends176 target-canvas units beyond each edge: target_xy =

↪ fullwide_display_xy - [176,176]. For a custom region, the exact display-to-target conversion

↪ appears in the tool result. Final answers always use the ORIGINAL target352 canvas, x/y0..351,

↪ upper-left origin.

Use additional batches to check a developing interpretation, earlier/later visibility, or trace

↪ continuity toward the target. You need not give coordinates for neighboring frames. If no

↪ issues are found, no temporal requests are needed. Never view the whole bank or images outside

↪ your requested batches.

PHASE C - FINAL REASSESSMENT.

Return sparse complete point replacements and sparse joint replacements. Unchanged records are

↪ preserved exactly. All13finalpoints and all6legs must remain present. You may revise any

↪ of13points when the audit identifies an issue; auxiliary joint changes are permitted for an

↪ audited leg. If new temporal evidence exposes another issue, describe it in a resolution with

↪ its node names rather than ignore it. Each audit issue must have a resolution: revised,

↪ retained, or unresolved, with supporting frame offsets and a short evidence statement. A

↪ provisional best-guess move can still be unresolved. A retained estimate can be well

↪ supported; an unresolved estimate must not acquire false certainty. End with final_check

↪ describing your visual check that the complete proposed pose is anatomically coherent,

↪ including unchanged landmarks and any remaining conflict. Do this check using already-viewed

↪ raw images and the assembled coordinates. No extra VLM calls, separate agents, or post-answer

↪ retry/overlay cycle.

ANATOMY: Three pairs of legs attach anteriorly (fore), centrally (middle), posteriorly (hind) to

↪ the thorax. Each chain is body -> coxa -> small trochanter/proximal femur -> femur -> tibia ->

↪ flexible five-part tarsus -> terminal foot. Proximal joints reorient, lift, advance and

↪ retract; the femur-tibia knee flexes; the tibia-tarsus and tarsal segments bend. The terminal

↪ leg is not rigidly collinear with the tibia. Projected lengths/angles vary with 3D

↪ articulation and foreshortening; legs can fold under body/wings, cross the midline and overlap

↪ each other. No fixed2Dangles, mirrored gait or imposed walking phase. Trace identity from

↪ attachment through crossings. Use visible portions to constrain hidden continuations; an

↪ occluded foot is not necessarily at the last visible trace or occluder boundary.

↪ Antennae/aristae originate on the head; wing veins move with translucent wings hinged on the

↪ thorax. Wings can open asymmetrically and the head/abdomen may bend. Hairs, shadows, walls and

↪ debris are not appendages.

LANDMARKS: head=head capsule center; thorax=thorax center; abdomen=posterior-most abdominal BODY

↪ tip (notcenter/wing); eyeL/eyeR=compound-eye centers; wingL/wingR=distal wing tips;

↪ forelegL4/forelegR4/midlegL4/midlegR4/hindlegL4/hindlegR4=terminal tarsal/foot tips. The

↪ target and temporal views may be dorsal, ventral, lateral or oblique; establish viewpoint from

↪ visible anatomy rather than imposing a dorsal template. Anatomical left/right belongs to the

↪ fly: when the head points up, its left is image-left in a dorsal view and image-right in a

↪ ventral view; rotate that convention with the animal. In lateral or oblique views, use

↪ attachment, body and wing relationships to distinguish visible near-side anatomy from occluded

↪ far-side anatomy. Image-left/right or near/far alone does not establish anatomical left/right.

↪ A viewpoint change across frames can clarify side identity; follow the same appendage back to

↪ the target without reflecting or copying a neighboring pose. If viewpoint or side remains

↪ ambiguous, preserve that ambiguity in the evidence and point identity confidence. Do not

↪ require projected symmetry or spread hidden legs into a dorsal template. Auxiliary joints:

↪ body_coxa=thoracic attachment; proximal_femur=femur base; femur_tibia=knee;

↪ tibia_tarsus=tarsus base.

For replacement points return xy(integer pair0..351), state(observed/inferred/prior_only),

↪ location_confidence and identity_confidence(high/medium/low),

↪ uncertainty_radius_px(integer1..176), reason(short). observed means a candidate is directly

↪ visible IN THE TARGET; a hidden target point supported by neighbors is inferred. prior_only

↪ means little image evidence constrains that particular location. These confidence/radius

↪ values are uncalibrated, and forced guesses need low confidence. Joint replacements have xy,

↪ state, confidence. Do not invent certainty or force coordinate changes merely to show activity.

No human labels, other flies’ outputs, earlier temporal-review results, web, models,

↪ segmentation/contours/edges/optical-flow algorithms, or coordinate fitting. Tools are

↪ restricted to supplied packet/target/overlay, your own audit/request/output files, and the

↪ supplied view tool plus the exact images it returns. No Vault or broader project reads;

↪ isolated context is necessary for this experiment. Save one response.json after the audit and

↪ bounded active investigation.

### C.4 Structured records

Initial responses contained an anatomical description, viewpoint and overlap assessments, 13 landmark records and six auxiliary leg chains. Point records specified integer coordinates in [0, 351], evidence state, location and identity confidence, uncertainty radius and supporting evidence. Uncertainty radii were integers in [1, 176] on the display canvas.

Audits specified landmark checks and questions for temporal review. Requests identified frame offsets, spatial regions and audit issues. Review responses supplied replacement records and issue resolutions. The image-access helper recorded coordinate transformations and which images had been displayed. Exact response schemas and the experimental template files accompany the manuscript source.

## References

[1] Gordon J. Berman, Daniel M. Choi, William Bialek, and Joshua W. Shaevitz. Mapping the stereotyped behaviour of freely-moving fruit flies. Journal of the Royal Society Interface, 11(99):20140672, 2014. doi: 10.1098/rsif.2014.0672. URL.

[2] Gordon J. Berman, William Bialek, and Joshua W. Shaevitz. Predictability and hierarchy in Drosophila behavior. Proceedings of the National Academy of Sciences, 113(42):11943–11948, 2016. doi: 10.1073/pnas.1607601113. URL.

[3] Talmo D. Pereira, Joshua W. Shaevitz, and Mala Murthy. Quantifying behavior to understand the brain. Nature Neuroscience, 23(12):1537–1549, 2020. doi: 10.1038/s41593-020-00734-z. URL.

[4] Alexander Mathis, Pranav Mamidanna, Kevin M. Cury, Taiga Abe, Venkatesh N. Murthy, Mackenzie Weygandt Mathis, and Matthias Bethge. DeepLabCut: markerless pose estimation of user-defined body parts with deep learning. Nature Neuroscience, 21:1281–1289, 2018. doi: 10.1038/s41593-018-0209-y.

[5] Talmo D. Pereira, Diego E. Aldarondo, Lindsay Willmore, Mikhail Kislin, Samuel S.-H. Wang, Mala Murthy, and Joshua W. Shaevitz. Fast animal pose estimation using deep neural networks. Nature Methods, 16(1):117–125, 2019. doi: 10.1038/s41592-018-0234-5. URL.

[6] Jacob M. Graving, Daniel Chae, Hemal Naik, Liang Li, Benjamin Koger, Blair R. Costelloe, and Iain D. Couzin. DeepPoseKit, a software toolkit for fast and robust animal pose estimation using deep learning. eLife, 8:e47994, 2019. doi: 10.7554/eLife.47994.

[7] Talmo D. Pereira, Nathaniel Tabris, Arie Matsliah, David M. Turner, Junyu Li, Shruthi Ravindranath, Eleni S. Papadoyannis, Edna Normand, David S. Deutsch, Z. Yan Wang, Grace C. McKenzie-Smith, Catalin C. Mitelut, Marielisa Diez Castro, John D’Uva, Mikhail Kislin, Dan H. Sanes, Sarah D. Kocher, Samuel S.-H. Wang, Annegret L. Falkner, Joshua W. Shaevitz, and Mala Murthy. SLEAP: A deep learning system for multi-animal pose tracking. Nature Methods, 19(4):486–495, 2022. doi: 10.1038/s41592-022-01426-1. URL.

[8] Anqi Wu, Estefany Kelly Buchanan, Matthew Whiteway, Michael Schartner, Guido Meijer, Jean-Paul Noel, Erica Rodriguez, Claire Everett, Amy Norovich, Evan Schaffer, Neeli Mishra, C. Daniel Salzman, Dora Angelaki, Andrés Bendesky, The International Brain Laboratory, John P. Cunningham, and Liam Paninski. Deep graph pose: a semi-supervised deep graphical model for improved animal pose tracking. In Advances in Neural Information Processing Systems, volume 33, pages 6040–6052, 2020. URL https://proceedings.neurips.cc/paper/2020/hash/4379cf00e1a95a97a33dac10ce454ca4-Abstract.html.

[9] Dan Biderman, Matthew R. Whiteway, Cole Hurwitz, Nicholas Greenspan, Robert S. Lee, Ankit Vishnubhotla, Richard Warren, Federico Pedraja, Dillon Noone, Michael M. Schartner, Julia M. Huntenburg, Anup Khanal, Guido T. Meijer, Jean-Paul Noel, Alejandro Pan-Vazquez, Karolina Z. Socha, Anne E. Urai, The International Brain Laboratory, John P. Cunningham, Nathaniel B. Sawtell, and Liam Paninski. Lightning pose: improved animal pose estimation via semi-supervised learning, Bayesian ensembling and cloud-native open-source tools. Nature Methods, 21:1316–1328, 2024. doi: 10.1038/s41592-024-02319-1.

[10] Chen Li and Gim Hee Lee. ScarceNet: Animal pose estimation with scarce annotations. In Proceedings of the IEEE/CVF Conference on Computer Vision and Pattern Recognition, pages 17174–17183, 2023. URL https://arxiv.org/abs/2303.15023.

[11] Shaokai Ye, Anastasiia Filippova, Jessy Lauer, Steffen Schneider, Maxime Vidal, Tian Qiu, Alexander Mathis, and Mackenzie Weygandt Mathis. SuperAnimal pretrained pose estimation models for behavioral analysis. Nature Communications, 15:5165, 2024. doi: 10.1038/s41467-024-48792-2.

[12] Tomas Jakab, Ankush Gupta, Hakan Bilen, and Andrea Vedaldi. Unsupervised learning of object landmarks through conditional image generation. In Advances in Neural Information Processing Systems, volume 31, 2018. URL https://arxiv.org/abs/1806.07823.

[13] Tejas D. Kulkarni, Ankush Gupta, Catalin Ionescu, Sebastian Borgeaud, Malcolm Reynolds, Andrew Zisserman, and Volodymyr Mnih. Unsupervised learning of object keypoints for perception and control. In Advances in Neural Information Processing Systems, volume 32, 2019. URL https://arxiv.org/abs/1906.11883.

[14] Jennifer J. Sun, Serim Ryou, Roni H. Goldshmid, Brandon Weissbourd, John O. Dabiri, David J. Anderson, Ann Kennedy, Yisong Yue, and Pietro Perona. Self-supervised keypoint discovery in behavioral videos. In Proceedings of the IEEE/CVF Conference on Computer Vision and Pattern Recognition, pages 2171–2180, 2022. URL https://arxiv.org/abs/2112.05121.

[15] Jennifer J. Sun, Lili Karashchuk, Amil Dravid, Serim Ryou, Sonia Fereidooni, John C. Tuthill, Aggelos Katsaggelos, Bingni W. Brunton, Georgia Gkioxari, Ann Kennedy, Yisong Yue, and Pietro Perona. BKinD-3D: Self-supervised 3D keypoint discovery from multi-view videos. In Proceedings of the IEEE/CVF Conference on Computer Vision and Pattern Recognition, pages 9001–9010, 2023. URL https://arxiv.org/abs/2212.07401.

[16] Tomas Jakab, Ankush Gupta, Hakan Bilen, and Andrea Vedaldi. Self-supervised learning of interpretable keypoints from unlabelled videos. In Proceedings of the IEEE/CVF Conference on Computer Vision and Pattern Recognition, pages 8787–8797, 2020. URL https://arxiv.org/abs/1907.02055.

[17] Jiteng Mu, Weichao Qiu, Gregory Hager, and Alan Yuille. Learning from synthetic animals. In Proceedings of the IEEE/CVF Conference on Computer Vision and Pattern Recognition, 2020. URL https://arxiv.org/abs/1912.08265.

[18] Siyuan Li, Semih Günel, Mirela Ostrek, Pavan Ramdya, Pascal Fua, and Helge Rhodin. Deformation-aware unpaired image translation for pose estimation on laboratory animals. In Proceedings of the IEEE/CVF Conference on Computer Vision and Pattern Recognition, 2020. URL https://arxiv.org/abs/2001.08601.

[19] Tomasz Niewiadomski, Anastasios Yiannakidis, Hanz Cuevas-Velasquez, Soubhik Sanyal, Michael J. Black, Silvia Zuffi, and Peter Kulits. Generative zoo. In Proceedings of the IEEE/CVF International Conference on Computer Vision, pages 8492–8502, 2025. URL https://arxiv.org/abs/2412.08101.

[20] Dongkai Wang, Shiyu Xuan, and Shiliang Zhang. LocLLM: Exploiting generalizable human keypoint localization via large language model. In Proceedings of the IEEE/CVF Conference on Computer Vision and Pattern Recognition, pages 614–623, 2024. URL https://arxiv.org/abs/2406.04659.

[21] Jie Yang, Wang Zeng, Sheng Jin, Lumin Xu, Wentao Liu, Chen Qian, and Ruimao Zhang. KptLLM: Unveiling the power of large language model for keypoint comprehension. In Advances in Neural Information Processing Systems, volume 37, 2024. URL https://arxiv.org/abs/2411.01846.

[22] Junho Kim, Hyungjin Chung, and Byung-Hoon Kim. CapeLLM: Support-free category-agnostic pose estimation with multimodal large language models. In Proceedings of the IEEE/CVF International Conference on Computer Vision, pages 22889–22898, 2025. URL https://arxiv.org/abs/2411.06869.

[23] Shiyu Zhao, Zhixing Zhang, Samuel Schulter, Long Zhao, Vijay Kumar B.G, Anastasis Stathopoulos, Manmohan Chandraker, and Dimitris Metaxas. Exploiting unlabeled data with vision and language models for object detection. In European Conference on Computer Vision, 2022. URL https://arxiv.org/abs/2207.08954.

[24] Geoffrey Hinton, Oriol Vinyals, and Jeff Dean. Distilling the knowledge in a neural network. arXiv:1503.02531, 2015. URL https://arxiv.org/abs/1503.02531.

[25] Yilei Chen, Xuemei Xie, and Fu Li. Vision-language model guided pose knowledge mining for human pose estimation. Journal of Computational Design and Engineering, 12(9):32–45, 2025. doi: 10.1093/jcde/qwaf079.

[26] Jingyang Ke, Weihan Li, Amartya Pradhan, Jeffrey Markowitz, and Anqi Wu. BehaviorVLM: Unified finetuning-free behavioral understanding with vision-language reasoning. arXiv:2603.12176, 2026. URL https://arxiv.org/abs/2603.12176.

[27] Olaf Ronneberger, Philipp Fischer, and Thomas Brox. U-Net: Convolutional networks for biomedical image segmentation. In Medical Image Computing and Computer-Assisted Intervention, pages 234–241, 2015. doi: 10.1007/978-3-319-24574-4_28.

[28] Benjamin R. Bauchwitz and Mary Cummings. Task configuration impacts annotation quality and model training performance in crowdsourced image segmentation. In Proceedings of the IEEE/CVF Winter Conference on Applications of Computer Vision, pages 6646–6656, 2025. URL https://openaccess.thecvf.com/content/WACV2025/html/Bauchwitz_Task_Configuration_Impacts_Annotation_Quality_and_Model_Training_Performance_in_WACV_2025_paper.html.

[29] Curtis Northcutt, Anish Athalye, and Jonas Mueller. Pervasive label errors in test sets destabilize machine learning benchmarks. In *Proceedings of the Neural Information Processing Systems Track on Datasets and Benchmarks*, volume 1, 2021. URL https://datasets-benchmarks-proceedings.neurips.cc/paper/2021/hash/f2217062e9a397a1dca429e7d70bc6ca-Abstract-round1.html.

[30] Grace C. McKenzie-Smith, Scott W. Wolf, Julien F. Ayroles, and Joshua W. Shaevitz. Capturing continuous, long timescale behavioral changes in Drosophila melanogaster postural data. PLOS Computational Biology, 21(2):e1012753, 2025. doi: 10.1371/journal.pcbi.1012753. URL.

[31] OpenAI. GPT-6 Astra: model documentation, 2026. URL https://developers.openai.com/api/docs/models/gpt-6-astra. Accessed 8 September 2026.

[32] Diederik P. Kingma and Jimmy Ba. Adam: A method for stochastic optimization. In International Conference on Learning Representations, 2015. URL https://arxiv.org/abs/1412.6980.

[33] Qwen Team. Qwen3.8-27B: model card, 2026. URL https://huggingface.co/Qwen/Qwen3.8-27B. Accessed 8 September 2026.

